# Toward standards in clinical microbiome studies: comparison of three DNA extraction methods and two bioinformatic pipelines

**DOI:** 10.1101/751123

**Authors:** Q.R. Ducarmon, B.V.H. Hornung, A.R. Geelen, E.J. Kuijper, R.D. Zwittink

**Author notes:** Address correspondence to Quinten R. Ducarmon.

## Abstract

When studying the microbiome using next generation sequencing, DNA extraction method, sequencing procedures and bioinformatic processing are crucial to obtain reliable data. Method choice has been demonstrated to strongly affect the final biological interpretation. We assessed the performance of three DNA extraction methods and two bioinformatic pipelines for bacterial microbiota profiling through 16S rRNA gene amplicon sequencing, using positive and negative controls for DNA extraction and sequencing, and eight different types of high- or low-biomass samples. Performance was evaluated based on quality control passing, DNA yield, richness, diversity and compositional profiles. All DNA extraction methods retrieved the theoretical relative bacterial abundance with maximum three-fold change, although differences were seen between methods, and library preparation and sequencing induced little variation. Bioinformatic pipelines showed different results for estimating richness, but diversity and compositional profiles were comparable. DNA extraction methods were successful for feces and oral swabs and variation induced by DNA extraction methods was lower than inter-subject (biological) variation. For low-biomass samples, a mixture of genera present in negative controls and sample-specific genera, possibly representing biological signal, were observed. We conclude that the tested bioinformatic pipelines perform equally with pipeline-specific advantages and disadvantages. Two out of three extraction methods performed equally well, while one method was less accurate regarding retrieval of compositional profiles. Lastly, we demonstrate the importance of including negative controls when analyzing low bacterial biomass samples.

**IMPORTANCE:** Method choice throughout the workflow of a microbiome study, from sample collection to DNA extraction and sequencing procedures, can greatly affect results. This study evaluated three different DNA extraction methods and two bioinformatic pipelines by including positive and negative controls, and various biological specimens. By identifying an optimal combination of DNA extraction method and bioinformatic pipeline use, we hope to contribute to increased methodological consistency in microbiome studies. Our methods were not only applied to commonly studied samples for microbiota analysis, e.g. feces, but also for more rarely studied, low-biomass samples. Microbiota composition profiles of low-biomass samples (e.g. urine and tumor biopsies) were not always distinguishable from negative controls, or showed partial overlap, confirming the importance of including negative controls in microbiome studies, especially when low bacterial biomass is expected.

## INTRODUCTION

Humans constantly interact with microbes that are present in the environment and reside on or within the human body. Recently, the attention for microbes has shifted from an exclusive interest in the pathogenicity of specific microbes toward the potential beneficial role of the microbiota in human health (1). The gastrointestinal tract contains the highest number of microbes and has been the most extensively studied body site of all human microbial communities (2). However, many other body sites are inhabited by various microbes composing a specific microbiome, such as the oral region, skin and urogenital system. Microbial complexity varies between these niches, e.g. the healthy vaginal microbiota is mainly composed of a few *Lactobacillus* strains (3), while gut and skin microbiota are more diverse (3).

A limiting factor in current microbiome research is that comparison of various study results is often difficult due to the application of different methodologies and lack of appropriate controls. These differences can affect data outcomes and lead to variation as large as biological differences (4). Variation can be introduced throughout the entire workflow, from sample collection, storage and processing to data analysis (5–8). Recently, more attention has been devoted to standardizing the workflow of microbiome research. For instance, it was observed that DNA extraction has a large impact on obtained data (4, 9) and consensus has been achieved regarding the application of bead-beating to increase efficiency of cell wall lysis and thereby improve the yield of Gram-positive bacterial DNA (10). Nevertheless, various kits and in-house extraction methods are used across different laboratories. Recently, Costea *et al.* evaluated 21 DNA extraction methods across three continents and suggested one protocol, named protocol Q, as ‘golden standard’ for human fecal samples. (9). They stated that it was unknown whether this method is optimal for other samples than fecal material, e.g. for low-biomass samples. To evaluate performance of DNA extraction for low-biomass samples, it is crucial to include multiple negative controls to allow for identification of bacterial DNA introduced during the entire workflow, from sample collection to sequencing (11).

As part of optimizing the procedures for 16S rRNA gene amplicon sequencing-based microbiome studies in our facility, we evaluated three DNA extraction methods and two bioinformatic pipelines using various positive controls and negative controls. In addition, we applied these DNA extraction methods to various biological specimens.

## MATERIALS AND METHODS

### Sample collection and pre-processing

Eight different biological specimens were included in this study, namely feces, urine, saliva, oral swabs, colorectal cancer tissue, colorectal cancer supernatant, vulvar squamous cell carcinoma tissue and formalin-fixed vulvar squamous cell carcinoma. Of each biological specimen, three unique samples were included. Only for oral swabs, six unique samples were included (Table S1). These samples were anonymized and treated according to the medical ethical guidelines described in the Code of Conduct for Proper Secondary Use of Human Tissue of the Dutch Federation of Biomedical Scientific Societies. A detailed overview of sample types, sample processing and storage conditions can be found in Table S1.

### Mock communities and DNA standard

Two mock communities (ZymoBiomics Microbial Community Standard, Zymo Research, Irvine, California, USA and 20 Strain Even Mix Whole Cell Material ATCC® MSA2002™, ATCC, Wesel, Germany) were included as positive controls for DNA extraction. Exact composition and relative abundances of 16S copies was provided on the product sheet for ZymoBiomics Microbial Community standard (hereafter referred to as Zymo mock), while for ATCC® MSA2002™ (hereafter referred to as ATCC mock) we calculated expected 16S profiles based on genomic information (Table S2). ZymoBiomics Microbial Community DNA Standard (hereafter referred to as DNA standard) was taken along as a positive sequencing control.

### DNA extraction

#### Procedures

Cancer samples were pre-processed for DNA extraction comparably to a recent study on pancreatic cancer microbiota (12), urine samples according to a recent publication on how to study urinary microbiota (13) and other samples according to in-house methods for sample processing (Table S1). For solid cancer samples, the beating steps during pre-processing were performed using a Qiagen TissueLyser LT (Qiagen Benelux, Venlo, the Netherlands) at 50Hz for one minute (Table S1). As single saliva samples did not contain sufficient volume for multiple extractions, several samples of the same individual were pooled to obtain the appropriate volume. DNA was extracted in duplicate from three unique samples for each biological material, only for oral swabs from six unique samples, and from the two mock communities. DNA was extracted using three different extraction protocols (see Protocols section), and for each protocol a negative extraction (no sample) was included in duplicate. The DNA standard was taken along in duplicate. DNA was quantified using a Qubit 3.0 Fluorometer (Invitrogen, Breda, the Netherlands) and the Qubit™ dsDNA HS Assay Kit (Thermo Fisher, Landsmeer, the Netherlands). A schematic overview of the study setup is shown in Figure 1.

**Figure 1:**
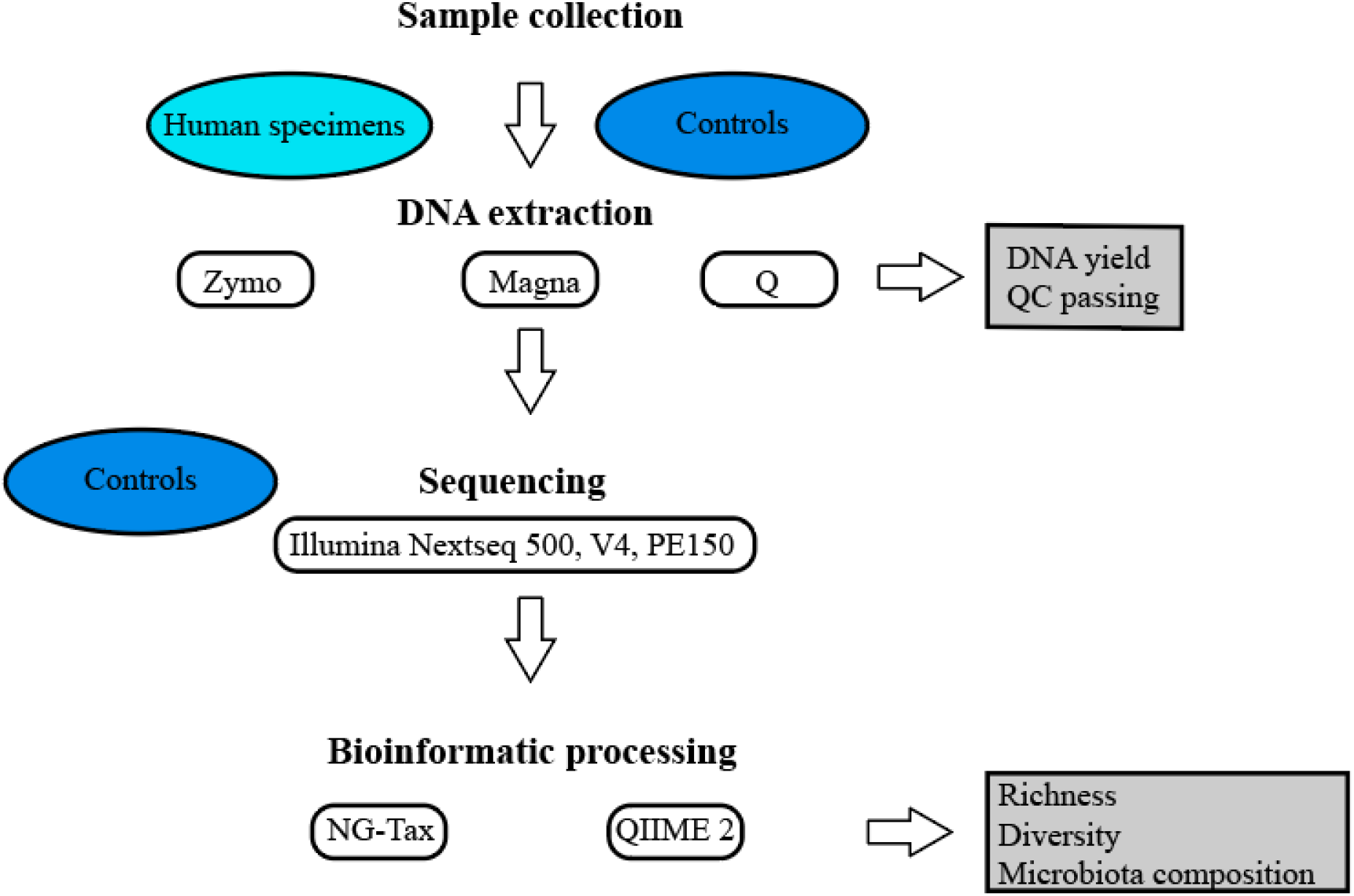
Study design workflow. DNA was extracted from human specimens and positive and negative controls using three different DNA extraction methods. DNA extraction performance was assessed on DNA yield and QC passing. Extracted DNA, and positive and negative sequencing controls were sequenced. Raw sequencing data was processed using two bioinformatic pipelines. Performance was assessed on microbiota composition, richness and diversity.

### DNA extraction protocols

Detailed protocols, including all minor adaptations, are present in Supplementary Methods. DNA extraction was performed using three methods: 1) the Quick-DNA Fecal/Soil Microbe kit (hereafter referred to as Zymo) (Zymo Research) according to manufacturer instructions with minor adaptations, 2) protocol Q (hereafter referred to as Q) (9) and 3) automated DNA extraction with MagNA Pure 96 ™ (hereafter referred to as Magna) (Roche Diagnostics, Almere, the Netherlands) using the MagNA Pure 96 DNA and viral NA small volume kit (Roche Diagnostics), according to standard operating procedures with minor adaptations. Mock communities were diluted to 10^4^-10^5^ cells per sample for extraction using Magna. For Q, several buffers and other materials were not provided in the kit and therefore purchased elsewhere, namely BeadBug™ prefilled tubes with 2.0 mL capacity and 0.1 mm Zirconium beads (Sigma-Aldrich, Zwijndrecht, the Netherlands), RNase A, DNase and protease-free water (10 mg/mL) (Thermo Fisher, the Netherlands) and TE buffer (Thermo Fisher).

### MALDI-TOF Mass Spectrometry (Biotyper)

To verify whether all bacteria of the ATCC mock were lysed after the first mechanical lysis step of both Zymo and Q, the lysate was plated on Blood Agar Plate, 5% Sheep Blood in Tryptic Soy Agar (VWR International, Amsterdam, the Netherlands) and aerobically and anaerobically incubated at 37°C for five days. The MALDI Biotyper system was used (Bruker Daltonics, Germany) to identify the bacterial species. Samples were prepared in the following way: A bacterial colony was taken from the culturing plate and spread in duplicate on single spots on a Bruker polished steel targetplate. Subsequently, one μl of 70% formic acid was added on each single spot and when dried, one μl prepared Bruker Matrix HCCA according to clinical laboratory protocols was added per spot. The Bruker polished steel targetplate was then used for MALDI-TOF MS Biotyper analysis.

### Library preparation and 16S rRNA gene amplicon sequencing

Of each duplicate DNA extraction from biological specimens, the duplicate with highest genomic DNA concentration was used for sequencing. Duplicate samples from controls were both sequenced. Quality control, library preparation and sequencing were performed by GenomeScan B.V. (Leiden, The Netherlands) using the NEXTflex™ 16S V4 Amplicon-Seq Kit (BiooScientific, TX, USA) and Illumina NextSeq 500 (paired-end, 150bp) according to their standard operating procedures. QC passing was based on intact genomic DNA and DNA concentrations measured by GenomeScan B.V. Therefore, those DNA concentrations were used for downstream analysis. Several samples were sequenced on multiple lanes, which is indicated in all relevant figures and tables.

### Sequencing data analysis

Read filtering, operational taxonomic unit (OTU)-picking and taxonomic assignment were performed using two different bioinformatic pipelines, QIIME 2 and NG-Tax 0.4 (14, 15), both using the Silva_132_SSU Ref database for taxonomic classification (16). The following settings were applied for QIIME 2: forward and reverse read length of 120, quality control using Deblur, identity level of 100%. A read length of 120 was chosen due to low quality sequence regions at the end of the reads. The following settings were applied for NG-Tax: forward and reverse read length of 120, ratio OTU abundance of 2.0, classify ratio of 0.9, minimum threshold of 1*10-7, identity level of 100%, error correction of 98.5. Prior to the NG-Tax run, potential left over primers were removed with cutadapt v. 1.9.1 (17), in paired-end mode, with additional setting −e 0.2 (increased error tolerance, 20%). This setting was required since database truncating based on the applied primers is part of the pipeline and, as such, primer sequences need to be removed to avoid mismatching with the database. Furthermore, all sequences with any deviating barcode in the fastq header were changed to the original barcode to allow inclusion into the NG-Tax pipeline.

The obtained OTU-tables were filtered for OTUs with a number of sequences less than 0.005% of the total number of sequences (18). Downstream analysis was performed in R (v3.5.1), mainly using the phyloseq (v.1.24.2) microbiome (v.1.2.1) and ggplot2 (v.3.0.0) packages (19–21).

### Data accessibility

All raw sequencing data used in the current study are deposited in the European Nucleotide Archive with accession number PRJEB34118.

## RESULTS AND DISCUSSION

### Mock communities pass quality control

We evaluated three different DNA extraction methods and two bioinformatic pipelines for microbiota profiling through 16S rRNA gene amplicon sequencing (Fig 1) using several positive and negative controls. Included positive controls were two bacterial mock communities and one DNA standard. Included negative controls were DNA extraction controls and sequencing controls. Quality control (QC) passing (DNA concentration and intact genomic fragment) were evaluated to determine extraction method performance. It was expected that positive controls would pass QC, while negative controls would not. Regarding mock communities, all extractions using Zymo and Q passed QC, while for Magna one extraction did not pass QC for both the ATCC mock community and Zymo mock community (Table S3). This was not unexpected, as mock communities were diluted for extraction using Magna and, therefore, DNA concentrations were lower. Negative extraction controls did not pass QC for Q and Magna, but they did for Zymo. This likely represents a higher contamination load during the extraction process for Zymo, which was also reflected by higher DNA concentrations (Table S3). A full overview of all samples included in this study, their QC passing and DNA concentrations can be found in Table S4.

### Positive controls: Classification, richness, diversity and relative species abundance

#### Primer choice may limit correct classification of all bacterial species in mock communities

Performance of the three extraction methods in combination with two bioinformatic pipelines, NG-Tax and QIIME 2, was evaluated on correctly identifying richness, diversity and relative abundances from bacterial mock communities and a DNA standard. Richness and diversity were computed at OTU level and at genus level. Analysis of compositional profiles was performed at genus level. Both pipelines failed to classify one organism from either mock community; NG-Tax did not detect *Cutibacterium* from the ATCC mock, while QIIME 2 did not detect *Salmonella* from the Zymo mock. The inability to detect *Cutibacterium* is most likely a primer choice issue, since the universal 515F and 806R primers are known to poorly amplify *Cutibacterium acnes* (22). This could be solved by choosing primers targeting different 16S regions, or by using adapted V4 region primers which do allow for accurate amplification of *Cutibacterium* (22, 23). Regarding QIIME 2 and the inability to detect *Salmonella*, there was an *Enterobacteriaceae* family with approximately expected relative abundance for *Salmonella*, and we were therefore confident this represented *Salmonella*. This *Enterobacteriaceae* family was subsequently included as *Salmonella*, and designated as *Enterobacteriaceae (Salmonella)*. This classification error likely resulted from the fact that *Enterobacteriaceae* members cannot always be discriminated based on the 16S rRNA V4 region (24).

#### DNA standard and Zymo mock community data can be recovered independent of extraction protocol or pipeline

The Zymo mock and DNA standard consist of respectively cell material or DNA of eight bacterial species and two fungal species. As the 16S rRNA gene was targeted, fungi should not be detected. Therefore, theoretical richness is eight and theoretical Shannon diversity was calculated to be 2.01.

Regarding the DNA standard, NG-Tax overestimated OTU-based estimated richness for both duplicates, DNA 1 and DNA 2 (Fig 2A, table S3). Richness was however accurately retrieved at genus level (Fig 2C). The same was observed regarding diversity, which was overestimated at OTU level (Fig 2B), but accurate at genus level (Fig 2D). QIIME 2 approached theoretical richness and diversity values at OTU level (Fig 2A+B, table S3). Richness estimates slightly improved at genus level (Fig 2C), while diversity did not differ from OTU-based diversity (Fig 2D). Thus, QIIME 2 better estimated richness and diversity at OTU level, while NG-Tax performed better at genus level (Table S3).

**Figure 2:**
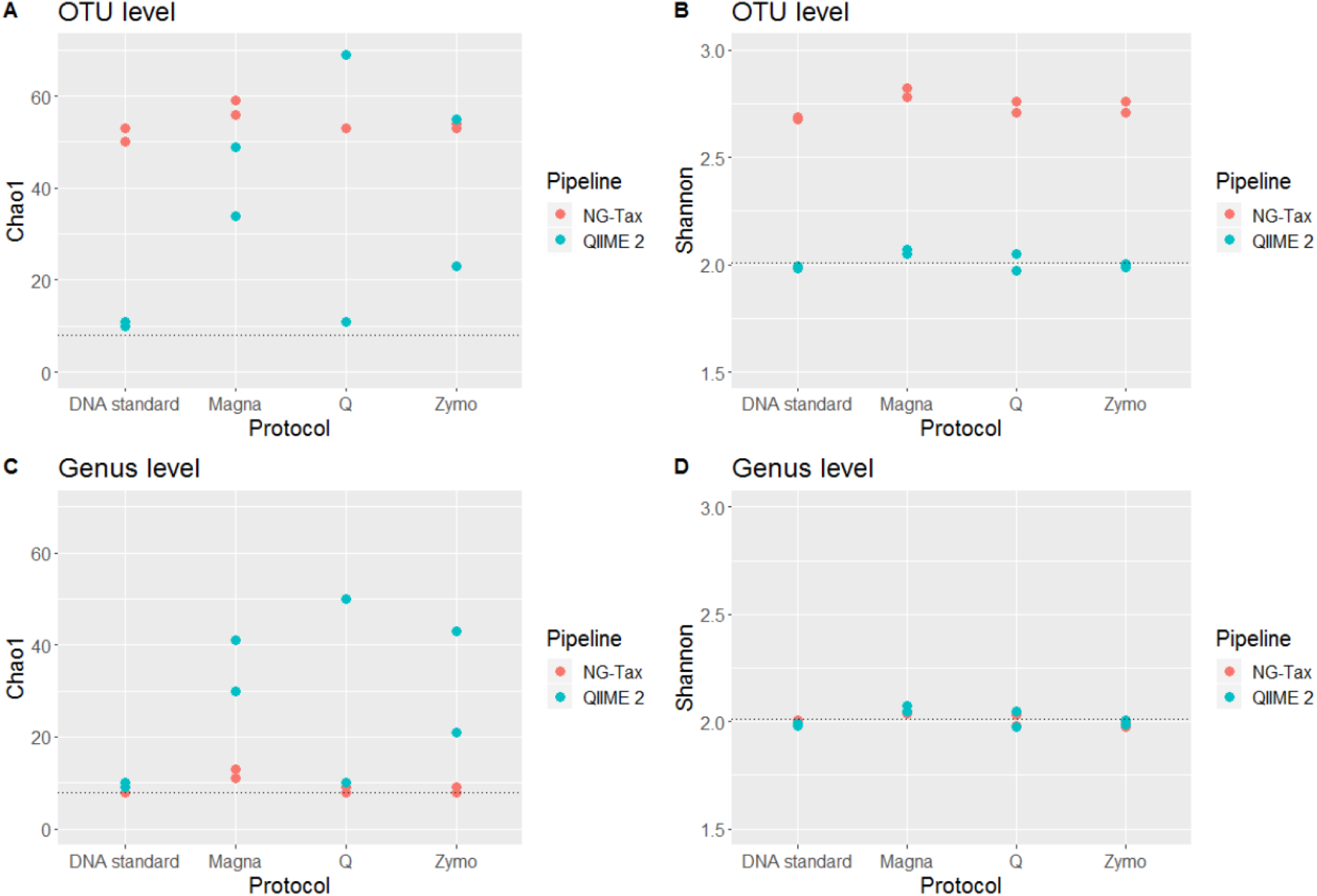
Richness (Chao1) and diversity (Shannon) computed for Zymo DNA and Zymo mock at OTU level (A+B) and at genus level (C+D) for each combination of bioinformatic pipeline and DNA extraction method. Dashed lines indicate theoretical values.

Compositional profiles of DNA 1 and DNA 2 are highly similar to theoretical abundance (Fig 3). To quantify differences in compositional profiles, Bray-Curtis dissimilarity and Kullback-Leibler divergence (Fig 4) (25) and fold errors for each taxon (Fig 5) were determined. For the dissimilarity and divergence values, a value of zero represents an identical microbiota composition to the theoretical expectation. NG-Tax obtained values closer to zero than QIIME 2 for both DNA 1 and DNA 2, although the difference is minimal (Fig 4 and Table S2) and the performance of both pipelines can therefore be regarded as equal. A similar conclusion can be drawn from the fold errors (Fig 5), since both pipelines accurately retrieved expected relative abundance, with all genera having a fold error between −1.5 and 1.5 (Table S3).

**Figure 3:**
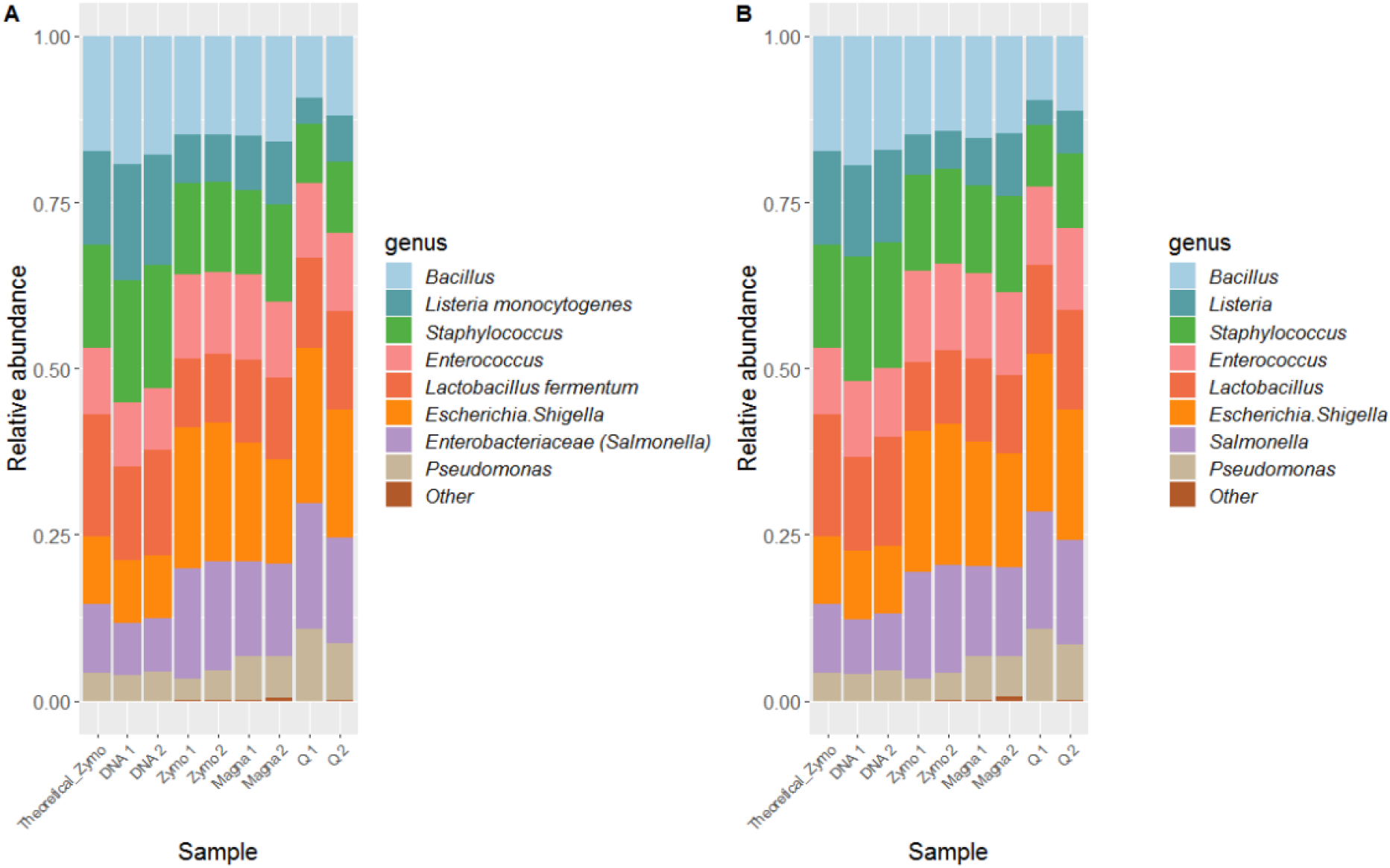
Compositional profiles at genus level for QIIME 2 (A) and NG-Tax (B) for Zymo mock, theoretical composition is indicated in the first bar graph.

**Figure 4:**
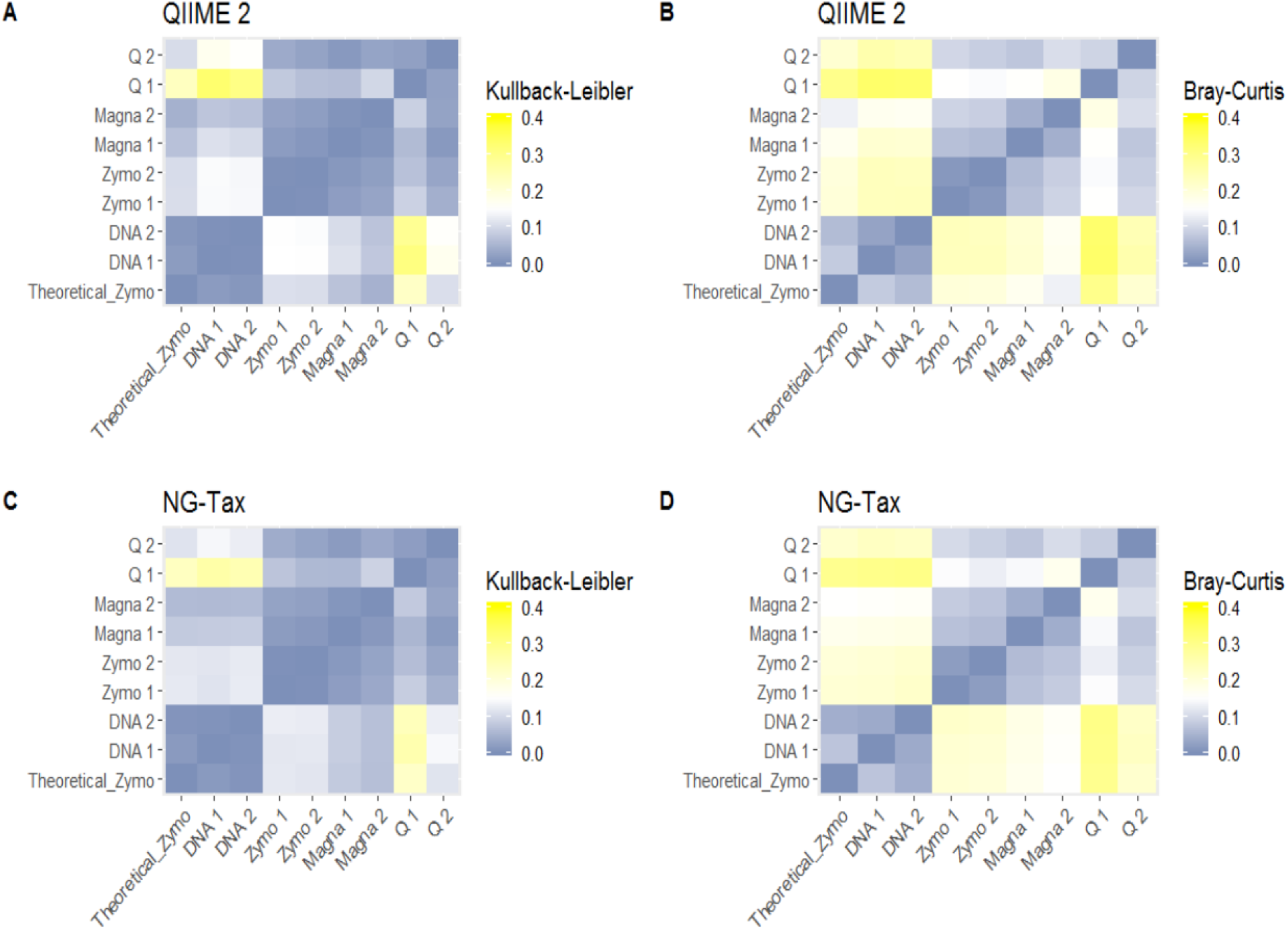
Comparison of compositional profiles expressed by Kullback-Leibler divergence (A+C) and Bray-Curtis dissimilarity (B+D) per pipeline. QIIME 2 results are shown in figure A+B, NG-Tax results are shown in figure C+D. For both Kullback-Leibler and Bray-Curtis, 0 indicates an identical compositional profile, while higher numbers indicate more dissimilar profiles.

**Figure 5:**
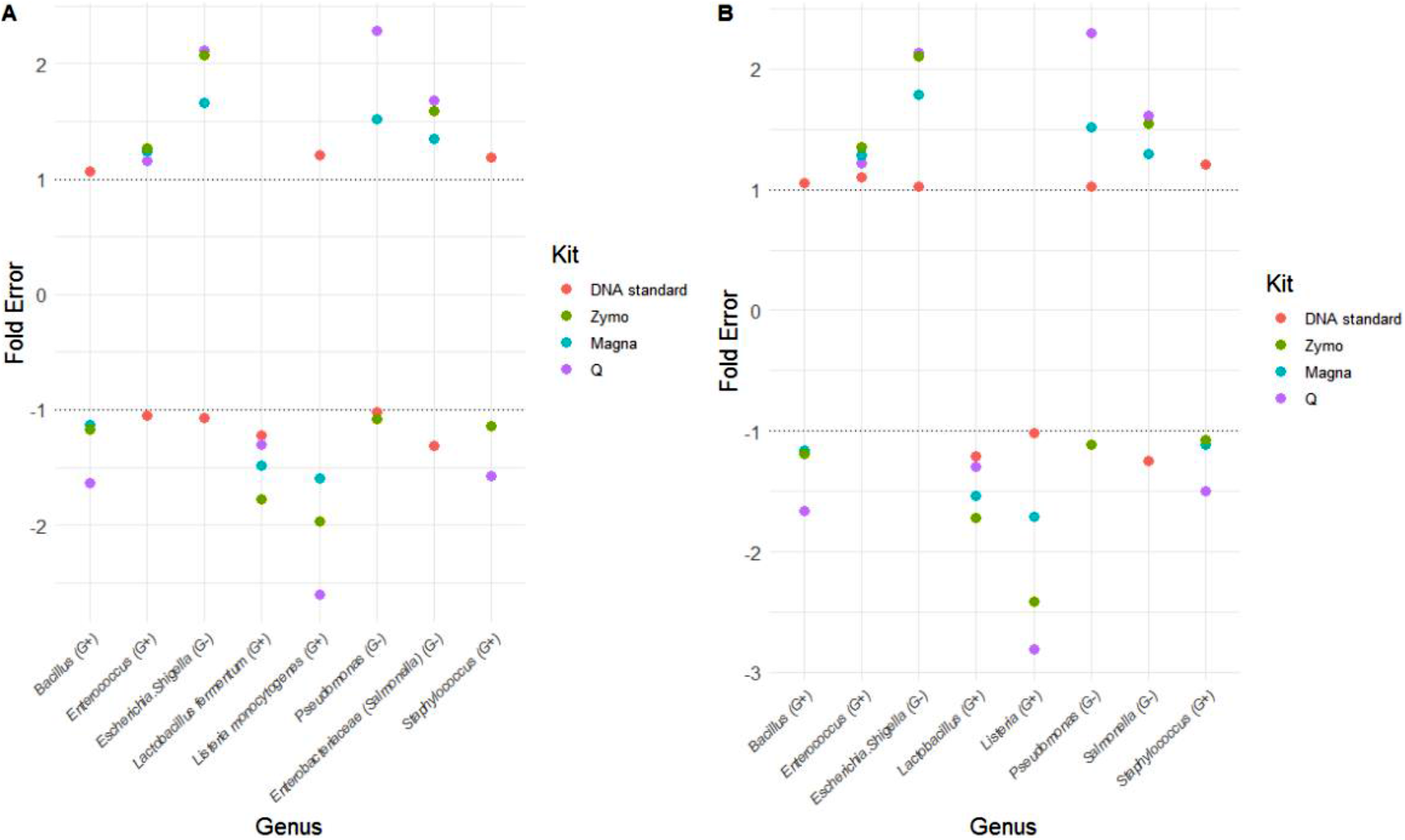
Fold error per bacterium as compared to theoretical values for QIIME 2 (A) and NG-Tax (B). A value above 1 represents overestimation, and a value below −1 represents underestimation.

Similar analyses were performed for the Zymo mock to evaluate performance of DNA extraction methods in combination with the bioinformatic pipelines. All DNA extraction methods, independent of pipeline, resulted in OTU-based richness above 20 for most samples, far higher than theoretical expectance (Fig 2A). This is especially noteworthy for QIIME 2, as it was highly accurate in retrieving correct richness for the DNA standard, in contrast to NG-Tax. Zymo and Q protocols in combination with NG-Tax retrieved accurate genus level-based richness, while a slightly inflated richness was observed for Magna (Fig 2C). No extraction method was consistent in retrieving correct genus level-based richness in combination with QIIME 2. Regarding diversity, all DNA extractions, independent of pipeline, retrieved highly accurate values at genus level (Table S3). At OTU-level, however, the NG-Tax pipeline resulted in overestimation of diversity independent of DNA extraction method, and can therefore be considered a result of bioinformatic processing. Magna extraction resulted in Bray-Curtis and Kullback-Leibler values closer to zero than Zymo and Q, independent of pipeline (Fig 4 and Table S3). A similar conclusion can be drawn from the fold errors, which are lowest for Magna and pipeline-independent (Fig 5 and Table S3). Taken together, results obtained from the DNA standard indicate that QIIME 2 and NG-Tax perform equally well in general, except for overestimation of OTU-level richness and diversity when using NG-Tax. Results obtained from the Zymo mock, which is a better representation of the full procedure for a microbiome study, indicate that richness is most accurate at genus level using protocol Zymo or Q in combination with the NG-Tax pipeline. In addition, bacterial microbiota composition profiles are best retrieved using Magna, followed by Zymo, and are pipeline-independent.

In concordance with current literature (9) and independent of extraction method, a general underestimation of Gram-positive bacteria was observed, with *Enterococcus* being the sole exception (Fig 5). This is most likely due to incomplete cell wall lysis of Gram-positive bacteria. Based on the DNA standard and the Zymo mock, we conclude that Zymo and Magna in combination with either pipeline are the best performing combinations (Table S3). However, when high-throughput DNA extraction is required (e.g. for large cohort studies), Magna may be preferred from a practical point of view, although it overestimates richness independent of pipeline.

In general, overestimation of OTUs may stem from the 100% identity setting for clustering, combined with the natural divergence of the 16S gene (26, 27). There is no current consensus on OTU identity setting, and cut-offs between 97% and 100% are used. An advantage of the 100% cut-off is that unique taxa differing a single nucleotide are clustered into different OTUs. A disadvantage is that, as intragenomic diversity in the 16S rRNA gene is common within bacterial genomes, a 100% cut-off can lead to multiple OTUs stemming from a single bacterium and thereby inflate richness (27). Apart from this biological explanation, the different algorithms and internal filtering steps used in QIIME 2 and NG-Tax can affect the outcome for richness.

#### ATCC mock is recovered incorrectly, independent of extraction protocol or pipeline

The ATCC mock consists of 20 unique bacterial species, with four of them belonging to two genera (*Staphylococcus* and *Streptococcus*). Therefore, theoretical richness at OTU level would be 20, but eighteen at genus level. In addition, these 20 unique bacterial species come from different environments, including gut, oral and skin microbiome.

No values close to the theoretical profiles for the ATCC mock for any extraction method/bioinformatic pipeline were observed, and one sample from Q consisted almost entirely of non-classifiable reads (Fig 6), indicating sample-related issues. *Bacillus* was highly overrepresented in all other samples, with a relative abundance over 30% in Zymo and Magna extracted samples, while 6.13% is expected. Curiously, after the first mechanical lysis step in Q, we could culture *Bacillus cereus* and *Cutibacterium acnes* (identification scores of 1.90 and 2.00, respectively), and *Bacillus cereus* (identification score 2.05) after mechanical lysis in Zymo. This is clinically important, as it means that infectious materials cannot be considered safe or non-infectious after mechanical lysis. As culturing of *B. cereus* indicates that cell wall lysis was incomplete, it would be expected that its relative abundance was underestimated, contrarily to what was observed. Another research group recently reported a similar overrepresentation of *Bacillus* in the ATCC community (28). ATCC itself was also unable to retrieve abundances close to theoretical expectation, neither with 16S amplicon sequencing nor with shotgun sequencing (29). Several reasons could explain this discrepancy between theoretical profiles and obtained profiles. For example, physical cell-to-cell interactions or presence of different metabolites may interfere with DNA extraction (26, 30). Therefore, based on this synthetic community, no conclusions on the optimal extraction-pipeline combination could be made. This proposed positive control prompts the question whether mock communities are always reliable for assessing performance of DNA extraction methods. As can be observed from the Zymo mock, DNA extraction kits do not necessarily inflict observed deviations, but may rather be a result of mock community-specific properties. Outcomes may depend on extraction kit / community type combination, indicating the potential necessity to use a positive control that strongly resembles the investigated microbiome.

**Figure 6:**
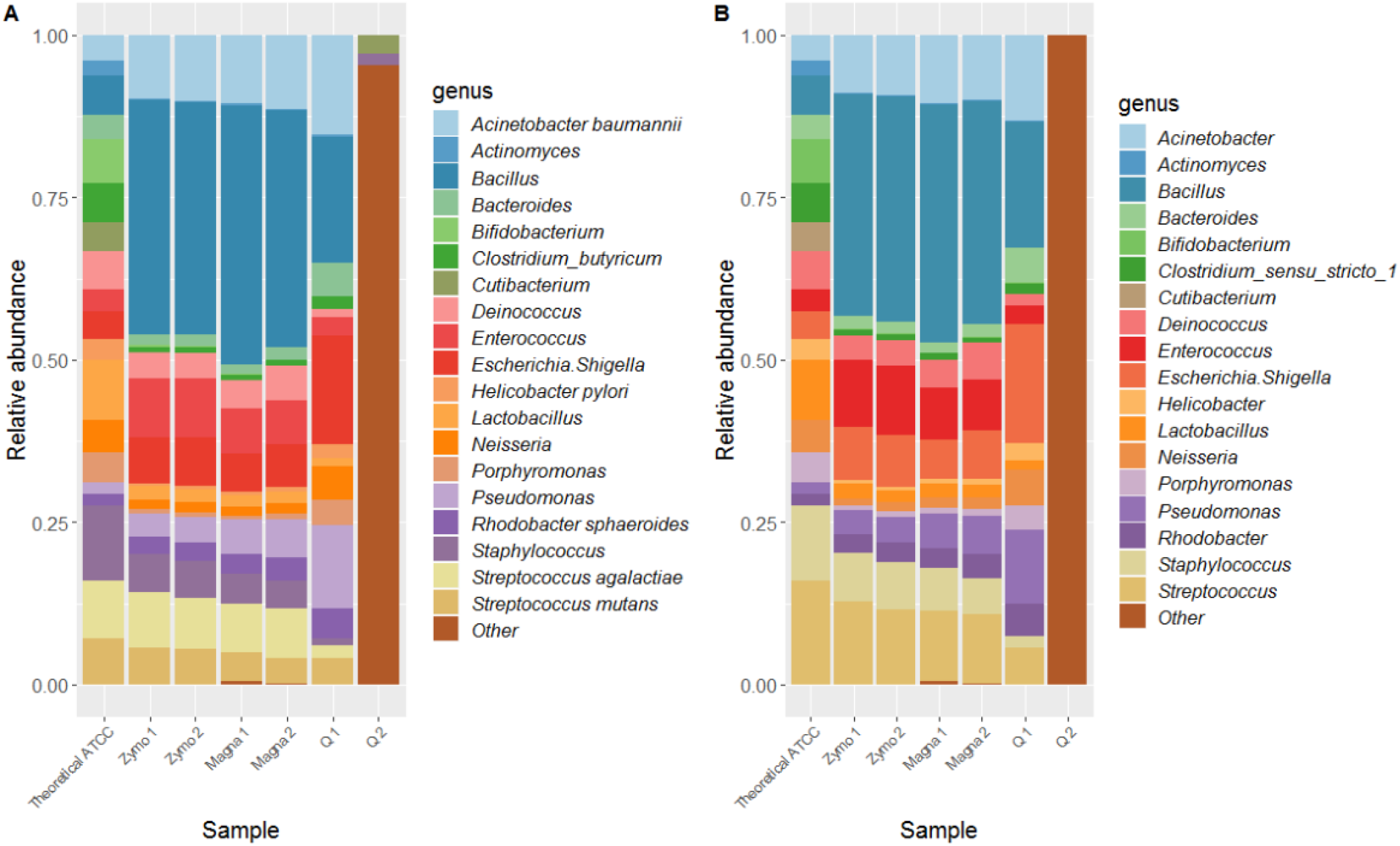
Compositional profiles at genus level for QIIME 2 (A) and NG-Tax (B) for the ATCC mock.

### Negative controls: Contaminating sequences are not always consistent

Negative controls were taken along for each extraction method to check for kit-specific contaminants, which is especially relevant for deciding whether low-biomass samples contain real microbiota. Regarding Zymo, clear kit-contaminants were *Pseudomonas* and *Delftia* (Fig S2A+C), consistent across the different pipelines at genus level, and with previous findings (11, 31). For Magna and Q, specific contaminants were less obvious, although *Pseudomonas* was present. Generally, negative controls mostly consisted of genera commonly found in gut and oral microbiota, most of them also previously described as contaminants (11). In addition, negative sequencing controls were taken along, and here no consistent contaminants could be observed (Fig S2B+D). Potential contamination sources are multifold, such as kit contamination, index hopping, or well-to-well contamination (32, 33). Index-hopping is however not a likely source of contamination, as the negative control for Magna was sequenced in different lanes, and profiles look highly similar (Fig S2A+C). Additionally, we did not observe index-hopping in our positive controls.

One of the contaminants we identified has not been previously described as a contaminant, namely *Clostridioides*. This likely represents *C. difficile*, and contamination by this bacterium can be explained by the fact that DNA extractions were performed in our National Reference Laboratory for *C. difficile*, which probably contains minor amounts of *C. difficile* spores during most time points. *C. difficile* contamination on laboratory surfaces has also recently been described in another clinical microbiology laboratory (34).

By incorporating this information with the Zymo positive controls, it can be concluded that Zymo and Magna are most optimal. Magna most accurately captured the expected community profile, while kit-specific contaminants are clear and easy to discriminate from biological signal using Zymo (Table S2). When investigating different biological sample types it might be warranted to use a kit for which kit contaminants do not overlap with the biological signal, e.g. *Pseudomonas* contamination when studying sputum samples from cystic fibrosis patients who are frequently colonized with *Pseudomonas* spp.

### Automatic Magna extraction yields lowest DNA for biological samples

Twenty-seven biological samples were available per extraction protocol (Table S1) and Q was most successful in passing QC (22/27), followed by Zymo (20/27) and Magna (17/27) (Table S3). DNA concentrations were on average lowest for Magna, while yields were comparable between Q and Zymo (Figure S1). Processing of raw sequencing data from biological samples was performed using the NG-Tax pipeline at genus level.

### The fecal microbiome can be sufficiently investigated independent of method

DNA extracted from fecal samples using the three different protocols all passed QC. Magna, Zymo and Q achieved an average concentration of approximately 29 ng/μl, 111 ng/μl and 212 ng/μl, respectively (Fig. S1). While DNA yield varied between extraction methods, all were sufficient for sequencing. Microbiota profiles were comparable between extraction methods for each sample (Figure S3A). In addition, differences in compositional profiles were quantified using Kullback-Leibler divergence (Figure 7A). This heatmap shows that technical variation induced by DNA extraction method is much lower than biological variation between feces samples. Profiles of the feces donors contained many bacterial genera commonly present in fecal microbiomes (35, 36). Healthy fecal microbiomes largely consist of Bacteroidetes and Firmicutes phyla (∼90%), while Actinobacteria and Proteobacteria are present in smaller proportions. At genus level, *Bacteroides*, *Prevotella* and *Faecalibacterium* are among the most prevalent genera, all of which were found in high abundance herein (3).

**Figure 7:**
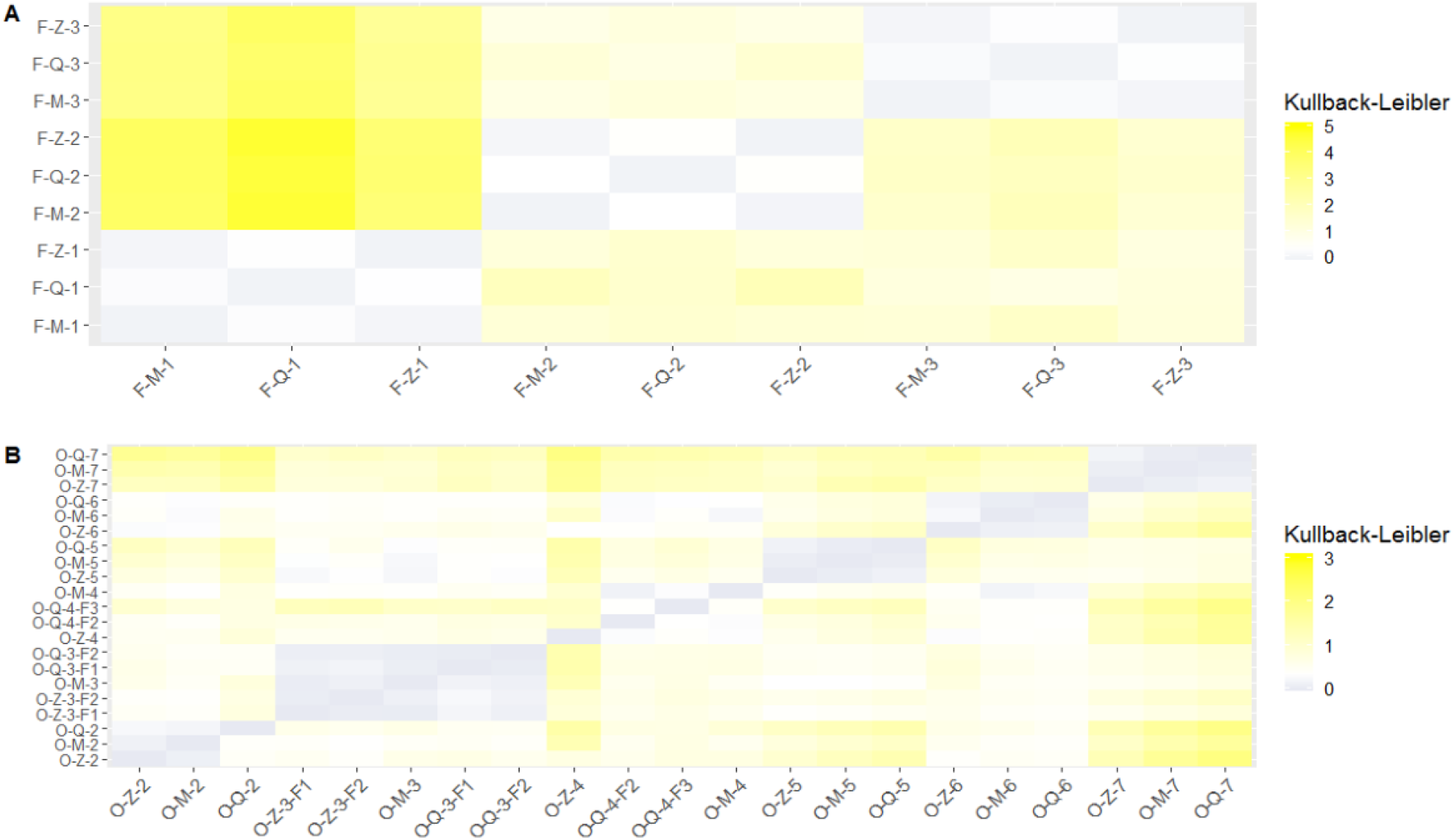
Kullback-Leibler divergence heatmap of feces (A) and oral swabs (B). Blue indicates highly similar composition, while yellow indicates divergence in composition. F1-F2-F3 represent samples which have been sequenced in duplicate, but on different flow cells.

### Low DNA yield of oral swabs does not seem to impact the microbiota profile

Out of eighteen DNA extractions, fifteen extractions passed QC for oral swabs. Only for Zymo, all extractions passed QC. DNA yields were highly variable for all extraction methods, ranging from 0.12 to 6.34 ng/μl. Half of the extractions (nine/eighteen) yielded a concentration below one ng/μl. All compositional profiles were dominated by *Streptococcus*, *Prevotella spp.*, *Haemophilus* and *Veillonella*, which was individual-independent. In addition, technical variation induced by DNA extraction and subsequent steps was lower than biological variation (Fig 7B). The oral microbiome, like the gut microbiome, is highly diverse. Nevertheless, a certain core of genera (e.g. *Streptococcus spp.* and *Prevotella spp.)* is present in most people, all of which were found in our study (3, 37, 38). Together, the good QC passing rate, DNA concentrations and consistency of compositional profiles between extraction methods lead us to conclude that all three methods work well for oral swabs.

### Applied methodology renders the urine microbiome unresolved

During the last decade, microbiome studies showed that urine contains a bacterial microbiota (39, 40). Despite using 30-40 ml of urine and centrifugation prior to extraction (13), we were not able to convincingly capture a urinary microbiota for all samples (Fig S3C). DNA concentrations were high for an infected sample (between thirteen and 42 ng/μl), but concentrations for the other samples were between 0.11 and 0.99 ng/μl. Six out of nine samples passed QC. For the infected sample with a high bacterial load, we were able to classify the cause of infection to *Enterobacteriaceae*, which is in agreement with the fact that most UTIs are caused by members of *Enterobacteriaceae*. One urine sample showed high similarity to negative controls for respective kits, with non-classifiable reads for Q and Magna, and high abundance of *Pseudomonas* for Zymo (Fig S3C). Another urine sample contained a high *Lactobacillus* abundance, which has previously been shown to be abundant in urine samples (40). In addition, presence of *Atopobium*, *Gardnerella*, *Campylobacter*, *Prevotella* and *Anaerococcus* point towards an existing urinary microbiota (41). However, *Pseudomonas*, a common Zymo kit contaminant, was still found in this urine sample, and for Magna more than 25% of reads could not be classified (Fig S3C). This could indicate that the biological signal is not much stronger than contamination, and therefore a mixed profile is observed. Further efforts and method optimization should be undertaken, although this can be difficult to implement in routine work (42). In addition, culturing could be used as a follow-up method to confirm that contaminants are not viable bacteria, but rather bacterial DNA.

### Applied procedures for saliva handling seem to be unsuitable for microbiome research

DNA yield from saliva samples was lower as compared to literature (43, 44) (Fig S1). Only a single DNA extraction had a concentration of slightly above one ng/μl (1.18; Table S4), while all other extractions had concentrations between 0.04 and 0.68 ng/μl. This may be associated with storage duration (∼fifteen years) and the fact that samples were thawed and refrozen several times. This also explains why only three out of nine DNA extracts passed QC. The included saliva samples were chosen as investigators within our facility were interested to see if microbiota studies could be performed using these samples.

Compositional profiles consisted of a mixture of genera present in the normal oral microbiota (*Oribacterium*, two *Prevotella* genera, *Streptococcus*, *Veillonella*) (3), genera present in our negative controls (*Pseudomonas*, *Delftia*) and non-classifiable reads (Fig S3D). In combination with low DNA yields, it is likely that a mixture between biological signal and contamination signal is present. Therefore, we consider the applied extraction methods unsuitable for saliva samples with a long duration of storage time and multiple freezing-thawing cycles.

### The colorectal cancer microbiome cannot be distinguished from negative controls or fecal microbiome

As colorectal cancer development has been associated with specific gut bacteria, we were interested to see if colorectal cancer tissue itself also contained bacteria (45, 46). DNA concentrations were sufficient for all samples to pass QC, but extracted DNA was likely mostly human-derived. Two of three extraction methods were not successful, as samples extracted using Zymo and Magna showed high similarity to their respective negative controls (Fig S3E). Using Q, *Bacteroides*, *Fusobacterium* and *Gemella* were identified, all being previously associated with colorectal cancer development (45, 47). Several gut commensals, including *Faecalibacterium* and *Escherichia-Shigella* were present in both the negative controls and these colorectal cancer samples. It is therefore difficult to discriminate whether these are contaminant bacteria, or whether they represent biological signal.

We hypothesized that by spinning down the material, the supernatant would contain more bacteria than the cancer tissue. DNA concentrations of supernatant were between 0.16 and 2.32 ng/μl, and seven out of nine DNA extractions passed QC (Table S4). For one sample, it was clear that across all methods many genera were observed which were present in negative controls (e.g. *Pseudomonas*), or reads could not be classified at all (Fig S3F). A second sample seemed to contain a real microbiota. Profiles were consistent across extraction methods, did not contain many contaminants and had specific bacteria previously linked to colorectal cancer (e.g. *Fusobacterium*) (45). The third sample showed a profile reflecting a mix between biological signal and technical contamination. Profiles were consistent across methods and contained genera representative of a gut microbiome, but also contained non-classifiable reads and contamination. Therefore, profiles are likely a mixture of biological signal and technical contamination, and further optimization is necessary prior to using this sample type for experimental studies. We have the same recommendation for colorectal cancer sample types as for urine, as discussed above.

### Vulvar squamous cell carcinoma does probably not contain bacterial DNA

Vulvar squamous cell carcinoma (VSCC) has different etiological pathways, of which one is associated with human papilloma virus (HPV). The counterpart is non-virally related and is frequently associated to lichen sclerosis, a benign chronic inflammatory lesion and *TP53* mutations (48, 49). We extracted DNA from HPV-negative VSCC tissue as a pilot study to determine if investigating the relationship between bacterial microbiota and HPV-negative VSCC would be potentially feasible. DNA concentrations were high (Fig S1), only for three extractions below one ng/μl, and eight out of nine extractions passed QC. However, DNA was probably again largely human-derived. This was reflected in the obtained microbiota profiles, as most reads were not classified or the profiles showed high similarity to negative controls (e.g. high abundance of *Pseudomonas*) (Fig S3G). Therefore, it is unlikely that this cancer tissue contains bacteria, or bacteria are so lowly abundant that they are overshadowed by contamination load. In general, the vulvar microbiome has not been extensively studied. A recent study on vulvar microbiome observed that *Lactobacillus*, *Corynebacterium*, *Finegoldia*, *Staphylococcus and Anaerococcus* are most abundant on this body site, but the use of negative controls was not reported (50). These genera are also part of the vaginal microbiota, and might be sampling contamination or reflect high similarity between vulvar and vaginal microbiota.

A large amount of formalin-fixed VSCC materials are stored in a biobank at our facility. To investigate whether this sample collection could be used for microbiota profiling, DNA was extracted from three formalin-fixed VSCC samples. DNA concentrations were all below 0.3 ng/μl, and only two out of nine extractions passed QC (Fig S4). One sample extracted with Q was excluded from further analysis, as no reads were present after sequencing. Extraction and sequencing of formalin-fixed material poses additional problems, as DNA molecules could be highly fragmented and too short for amplicon sequencing of the V4 region (51). For Zymo, samples resembled negative controls, with *Delftia* and *Pseudomonas* being highly abundant (Fig S3H). The same samples had completely different microbiota profiles when using protocol Q or Magna. Both extraction methods showed genera commonly found in the lower urogenital tract, including *Staphylococcus*, *Streptococcus*, *Prevotella* and *Gordonia* (3, 36). However, many of these genera were also detected in negative controls. In combination with low DNA yield and inconsistent profiles across extraction methods, we conclude that no reliable bacterial microbiota profile could be identified in these samples. For both VSCC types, we suggest the same way forward as for urine samples.

### Sample groups with and without biological signal cluster apart

Lastly, we performed t-distributed stochastic neighbor embedding (t-SNE) clustering using Bray-Curtis measures on all samples used in the present study (Fig 8) (52). Based on microbiota composition as measured by Bray-Curtis, t-SNE projects points in a two-dimensional space, while maintaining local structures present in high-dimensional space. Clear clusters could be identified for Zymo positive controls, feces, oral swabs and ATCC mock (all but one sample) (Fig 8). Other biological samples and negative controls were more dispersed throughout the plot, indicating that either more biological or technical variation was present. This is in agreement with our detailed analysis, showing that their microbiota cannot necessarily be distinguished from the negative controls. An example of the importance of including negative controls comes from two studies aiming to unravel the placental microbiota (53, 54). It is currently unclear whether a placental microbiota exists, but when comparing placental samples of healthy deliveries to included negative controls, microbiota compositions could not be distinguished (53, 54).

**Figure 8:**
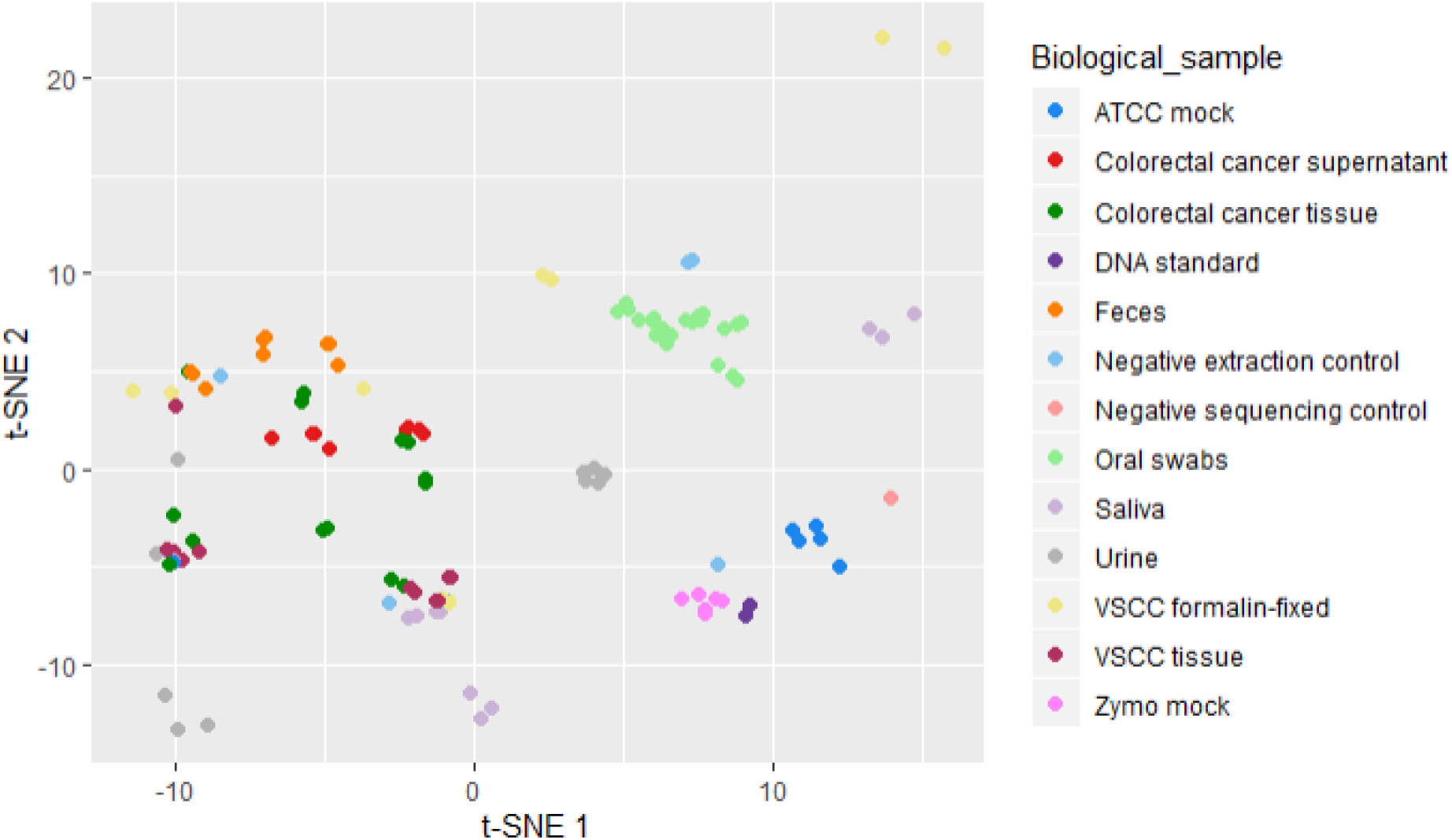
Bray-Curtis distance measures visualized by t-distributed stochastic neighbour embedding (t-SNE) for all samples. Each dot in the plot represents a single sample, and short distances between samples indicate high similarity.

### Strengths and limitations

The current study had several strengths and limitations. By using a positive control of cell material with a corresponding DNA standard, we differentiated variation induced from sequencing procedures and DNA extraction. We demonstrate the importance of using positive and negative controls in microbiome studies, and show that negative controls are crucial for interpretation of low-biomass samples. Another strength of the study was that for several biological samples (feces and oral swabs), we showed that technical variation was much smaller than biological variation. A shortcoming of the study is that we did not perform any other quantification next to 16S sequencing (e.g. qPCR), which may be particularly useful for quality control of the ATCC mock. Furthermore, the current study used only three unique samples of most biological sample types. Especially for samples for which DNA extraction was challenging (urine samples, colorectal cancer supernatant), a higher number of unique samples would have allowed for a more thorough evaluation.

## CONCLUSION

The current study evaluated three DNA extraction methods and two bioinformatic pipelines for bacterial microbiota profiling using several positive and negative controls, and a range of biological specimens. All three extraction methods quite accurately retrieved theoretical abundance of the Zymo mock, but not of the ATCC mock. For DNA extraction, we recommend using the Zymo and Magna protocol, since they showed good overall performance for all samples. Sequencing procedure only induced minor variation, as shown using a DNA standard. We furthermore showed that the NG-Tax and QIIME 2 pipelines perform equally well overall, each having their specific flaws.

By including negative controls and comparing these with low-biomass samples, we evaluated whether low-biomass samples consisted of technical noise, biological signal or a mixture. In most cases, identification of a unique microbiome was not achieved, highlighting the importance of negative controls and sufficiently sensitive methods. The results from this study can help other microbiome study groups to select an appropriate DNA extraction method and bioinformatic pipeline. We hope this study contributes to further standardization in methodology in the microbiome field, and to increased awareness of the usage of controls, especially when studying low-biomass samples.

## ACKNOWLEDGEMENTS

We thank all collaborating partners who provided us with clinical biospecimens, namely Liz Terveer, Eric Berssenbrugge, Erik Giltay, Noel de Miranda, Jitske van den Bulk, Natalja ter Haar and Kim Kortekaas. We also thank Eric Claas for permission for using the MagNA Pure 96 ™ and the clinical laboratory for identification of *Bacillus cereus* and *Cutibacterium acnes* using MALDI-TOF.

## AUTHOR CONTRIBUTIONS

QD, BH, AG, EK and RZ conceptualized and designed the study. QD and AG performed practical laboratory work. BH and RZ processed raw sequencing data. QD analyzed data, prepared figures and wrote the manuscript under supervision of BH and RZ. All authors interpreted data, read and revised drafts of the manuscript, and approved the final version.

